# MoMA: Large scale network model of Microbes, Metabolites and Aging hallmarks

**DOI:** 10.1101/2023.08.28.555041

**Authors:** Sarvesh Menon, Nishad Bapatdhar, Bipin Pradeep Kumar, Samik Ghosh

## Abstract

The gut microbiome is known to be a driver of age-related health decline. Various studies have shone light on the role of the gut microbiome as a marker as well as modulator of aging processes. However, the mechanisms by which the microbiome affects aging are still unclear. We have developed a Microbiome Metabolite Aging (MMA) fusion network by building upon a metabolic interaction network of gut microbiota to develop associations with the hallmarks of aging. The MMA, consisting of 238 metabolite-aging hallmark interactions serves as a tool to investigate the mammalian (and in particular human) gut microbiome as an effector of aging at a systems-level. The network further identifies 249 microbes that unequivocally affect the hallmarks of aging. The results highlight how the underlying biology of microbial metabolite mediated interactions, in conjunction with the topological properties at a network level, differentially regulate the aging hallmarks. This detailed microbial and metabolite association to the hallmarks of aging provides a foundation which is envisaged to be instrumental in advancing our knowledge of the physiology of aging, and for the development of novel therapeutic tools.

## 1 Introduction

Numerous biological associations and various health mediators can be found within the vast microbial colonization of the human gut. The estimates of microbes present in the human gut vary widely in different studies, but it is generally accepted that we harbor more than 1000 microbial species-level phylotypes (Rajilić-Stojanović and de Vos 2014). These microbes form a complex ecology that influences human physiology, susceptibility to illnesses and diseases, even how we age.

There are an increasing number of studies in aging research which focus on the contributory role of the gut microbiome as a modulator of aging (Ghosh, Shanahan, and O’Toole 2022; Kim and Benayoun 2020; Ragonnaud and Biragyn 2021; Badal et al. 2020). The research even identifies facets of host physiology that suggests a bidirectional relationship between the gut microbiome and aging (Kim and Benayoun 2020). where a direct causal role of the gut microbiome on host aging has been suggested by a number of studies using various experimental models. Studies have identified how the changes in gut microbiota across an individual’s lifespan are influenced by age-related processes. At the same time, association of gut microbiota with the hallmarks of aging (López-Otín et al. 2013; Kowald and Kirkwood 1996) have not been systematically mapped or modeled. Scientific understanding of the molecular underpinnings of aging related hallmarks have developed further in (López-Otín et al. 2023), where they build on their previous system with additions based on discoveries of the last decade.

The new system proposed the set of 12 Hallmarks of Aging as genomic instability, telomere attrition, epigenetic alterations, loss of proteostasis, disabled macroautophagy, deregulated nutrient-sensing, mitochondrial dysfunction, cellular senescence, stem cell exhaustion, altered intercellular communication, chronic inflammation, and dysbiosis. Hallmarks of aging were defined to fulfill three criteria: (1) The hallmark manifests over time, accompanying aging, (2) Accentuating the hallmark must accelerate aging, and (3) Interventions on the hallmark presents an opportunity to decelerate or reverse aging.

Metabolites are intermediate products of metabolic reactions. Microbiota-derived metabolites range from exogenous dietary substrates or endogenous host compounds and mediate a wide array of effects on our metabolism. Classes of microbiota-derived metabolites include bile acids, SCFA (Short-Chain Fatty Acids), BCAA (Branched-Chain Amino Acids), Trimethylamine N-oxide, Tryptophan and indole derivatives (Agus, Clément, and Sokol 2021). Martin et al (Martin et al. 2019) establish clear relationships between gut bacteria and host metabolism in which microbe-mediated gut hormone release plays an important role. In the context of aging, a set of core metabolites are emerging as central mediators of aging (Sharma and Ramanathan 2020). However, there is little known of microbial metabolites that are connected to the organized hallmarks of aging, or even aging in general.

In this work, we systematically collect, curate and harmonize the current state-of-the-art knowledge on the complex interaction dynamics between microbes, their metabolites and their connection to the hallmarks of aging. We build upon a metabolic interaction network of the mouse and human gut microbiota (Lim et al. 2020), comprising 838 microbial species interacting through 8,224 small-molecule transport and macromolecule degradation events to develop associations with the hallmarks of aging to construct the Microbiome Metabolite Aging (MMA) fusion network. The network captures directional and where available, causative interactions between metabolites and the hallmarks with a focus on the impact of the gut microbiome and associated metabolites on the hallmarks of aging.

As a broad framework connecting microbes to metabolites and then metabolites to the hallmarks of aging, conceptually illustrated in Fig.1, the MMA serves as a tool to investigate the mammalian (and particularly human) gut microbiome as an effector of aging at a systems-level. Using the MMA network model, we focused analysis on studying the effect of microbes and metabolites on the 12 hallmarks of aging, in terms of various network properties like degree, eigenvector centrality as well as path-length analysis. The results, as elucidated in later sections, highlight how the underlying biology of microbial metabolite mediated interactions, in conjunction with the topological properties at a network level, differentially regulate the aging hallmarks.

**Figure 1.**
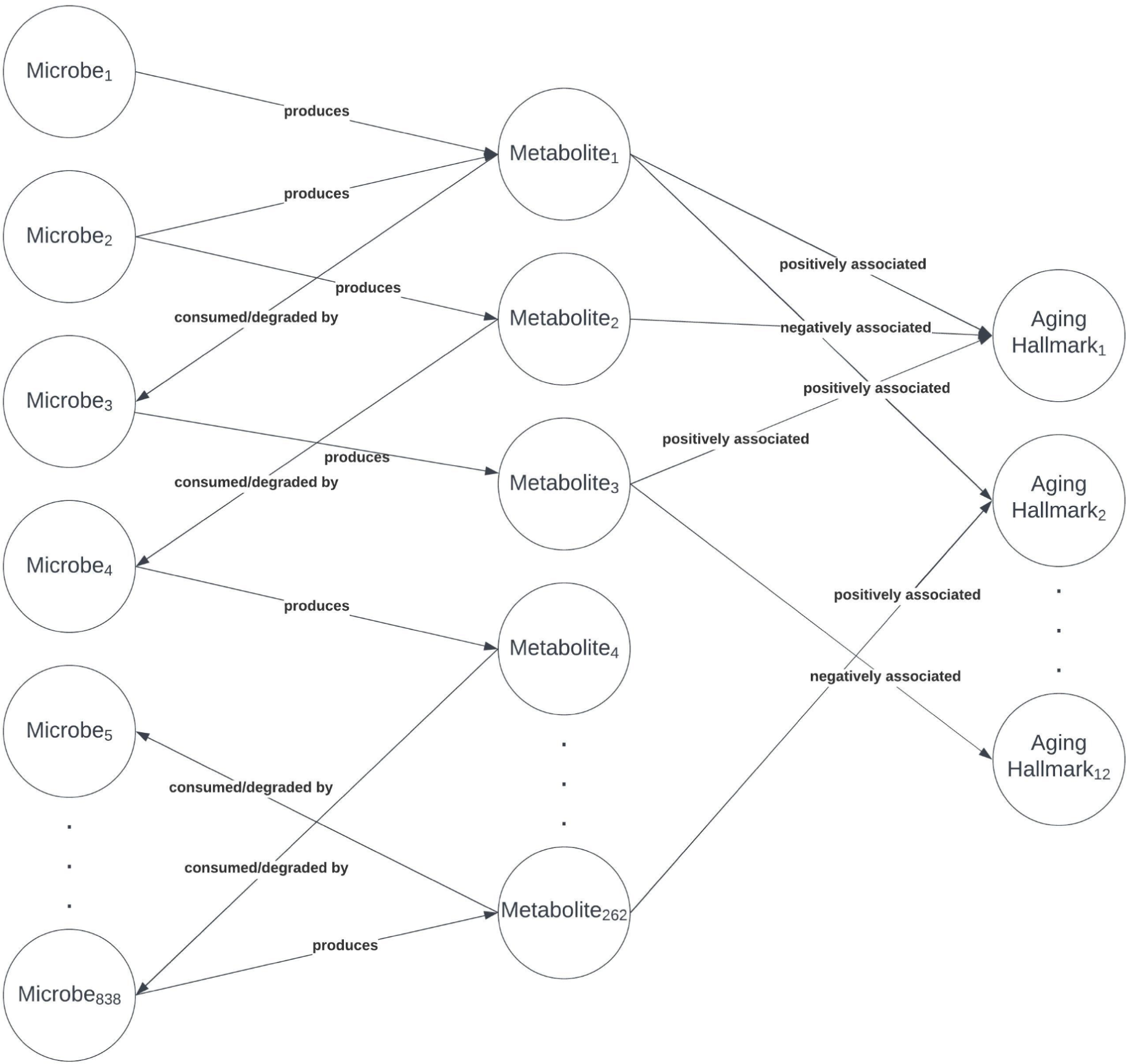
Schematic representation of MMA network, describing the types of nodes and edges present.

This detailed microbial and metabolite association to the hallmarks of aging provides a foundation which is envisaged to be instrumental in advancing our knowledge of the physiology of aging, and for the development of novel therapeutic tools.

## 2 Materials and Methods

### 2.1 Knowledge network creation

Curation of the MMA Network consisted of two parts: A microbiome-metabolite network and a metabolite-aging hallmark network, the steps involved in the generation of both of them are elucidated below.

#### 2.1.1 Microbiome - metabolite network

The first part of the MMA network, the microbiome-metabolite network, required a characterization of the stoichiometry of how microbes interact with metabolites. For this, we used the extensive literature-curated interspecies network of mammalian gut microbiota called NJC19 (Lim et al. 2020). The complete file was downloaded from the supplementary table associated with the publication. Following which, non-network data was removed (such as explanation lines and serial numbers), and only the species, small-molecule metabolite or macromolecule and metabolic activity column were retained. Negative associations (associations known to not exist), marked by a (-) in the metabolic activity were removed. Mouse tissue cells i.e., mouse goblet cells, mouse hepatocytes and mouse colonocytes and all their associations were removed from the network. Rows containing multiple associations were split into separate rows.

#### 2.1.2 Metabolite - aging hallmark network

To create a Metabolite-aging hallmark network, the list of metabolites to be reviewed was the set of unique metabolites in the microbiome-metabolite network. This gave us a list of 262 metabolites to analyze. Then the structure was determined to query existences of associations between the identified metabolites and the hallmarks. The protocol applied in this study is elucidated below, including the eligibility criteria, information sources, search strategy and query strings. Associations to aging hallmarks for humans were the primary target, but literature which stated the associations’ possible existence in humans by citing studies on model organisms *Mus musculus* and *C. elegans* were also used.

For each metabolite, the pubmed query was “[Metabolite Name]” AND (gut OR Humans) AND (Aging OR “[Hallmark Name]”). Where metabolite name and hallmark name (square brackets to mark variables and not in the actual query) were run across the table iteratively, for both hallmarks and metabolites, substrings of the whole string were also used as queries, for example “PABA” in the case of metabolite “4-Aminobenzoate (para-aminobenzoic acid, PABA, p-Aminobenzoate)”. Google scholar was also searched using the query “[Metabolite Name]” gut humans [Aging/Hallmark], where metabolite name follows the same rules above and aging/hallmark follows the same rules as Hallmark name but also included Aging in the list. Papers cited in identified papers were selected on a case-by-case basis to be reviewed for possible associations between hallmarks and metabolites.

Among identified papers, for hallmarks genomic instability (referred as DNA damage and genomic instability in the analysis), telomere attrition, disabled autophagy, mitochondrial dysfunction, cellular senescence, stem cell exhaustion, dysbiosis and chronic inflammation, only in the case of explicit reference to the hallmarks, string subsets of hallmarks (e.g., inflammation for chronic inflammation), synonymous words (e.g., macroautophagy in place of autophagy), or any combination of the same (e.g., Reduced stem cell proliferation for stem cell exhaustion), were the metabolites marked to be associated with the hallmarks.

In the cases of epigenetic modifications, histone modifications, post translational modifications and non-coding RNA interactions, which the research explicitly links to aging were used as the Hallmark epigenetic alterations.

For loss of proteostasis, protein homeostasis, any term associated with protein folding/misfolding explicitly or chaperonins, and protein production mechanism or transport functioning (explicitly under protein production or transport) was considered.

For nutrient sensing, dysregulation of IGF-1, mTOR or AMPK pathways were considered synonymous, disruption of glucose homeostasis and other pathways when mentioned in the context of nutrient sensing pathways linked to aging were also allowed (e.g., SIRT family-associated pathways).

In the case of dysbiosis, metabolites linked with increase of specific microbes which in turn are associated with dysbiosis were passed on, as effects of metabolites on the microbes are to be captured through the microbiome-metabolite network.

In cases of explicit reference to a sequential effect on multiple hallmarks of aging, such as a metabolite causing inflammation which in turn causes DNA damage, only the hallmark affected by the metabolite directly is added to the network. This step was taken in accordance with the network capturing the impact of the metabolites on the hallmarks independently. Furthermore, cases of explicit reference to terms delinking the effect from being an age-associated or chronic case were also removed, the prefix of acute, such as in acute inflammation, or short-term and temporary as prefixes.

The identified associations were then classified as either beneficial or detrimental to the hallmark of aging. In this context, a beneficial association between a given metabolite and a hallmark implies that the metabolite accelerates host aging through that specific hallmark.Positive associations were listed with a 1 and negative associations with a −1. This network was then represented as three columns; Metabolite, Hallmark, Interaction, where the direction of the association is understood to be from the metabolite to the hallmark.

Our research focused on identifying interactions between metabolites and specific hallmarks of aging. As a result, we did not consider the impact of metabolites on age-related conditions that are not explicitly linked to a hallmark or aging. This approach also led us to exclude 157 metabolites that were not directly implicated in the aging process through a hallmark but may have had indirect associations with age-related conditions, along with possible other interactions of the metabolites included. Addressing these indirect associations would require a thorough understanding of the interactions between the hallmarks and these age-related conditions, which falls outside the scope of our analysis.

### 2.2 Network processing

The edited NJC 19 network and the curated pairwise Metabolite-Hallmark association file were merged to give the universal MMA network file, with the columns labeled as Source, Target and Interaction. The network was processed from text-based information to quantitative (only ternary information as shown next) to analyze the topology of the network. To represent associations between microbes and metabolites, we replaced “Production (export)” interactions with a +1, and “Consumption (import)” and “Macromolecule degradation” interactions with a −1. This signifies the microbe’s impact on the metabolite, reflecting the change in metabolite levels caused by an increase in the microbe.

The table was converted into a network with the Networkx library in python(Hagberg, Swart, and S Chult 2008). Using the nx.add_weighted_edges_from function for a weighted graph (indicating the different types of associations) and nx.add_edges_from for an unweighted graph (for counting-associated analysis) and subsequent analysis was carried out using functions in the library. Weighted graphs were used to capture the ternary nature of the interactions, with +1, 0 and −1 weights, for node information metrics to differentiate types of interactions and the lack of an interaction. To capture the impact of the microbes on the metabolites, human host cell nodes were deleted from the network, the cells removed were the human goblet cells, human hepatocytes and human colonocytes, as the current network models direct interaction of the gut microbes with the metabolites present and the resultant equilibrium metabolic fluxes affects the host physiology.

#### 2.2.1 Directed network generation

A directed network graph was generated to model the transport of the metabolite in the gut, wherein metabolite nodes represented the metabolite in the environment. Following this, edges into the metabolite node imply the metabolite being released in the environment, conversely edges out of it imply the metabolite being removed from the environment. In the directed network graph, the processed MMA file was used with the Source and Target columns as the source and target nodes respectively. The negative associations between microbes and metabolites were formatted to swap the source and target columns to have the metabolite as the source and the microbe as the target, following the direction of the metabolite transport as shown in NJS 16(Sung et al. 2017). The directed network graph was generated using nx.Digraph and nodes and edges were loaded using the above functions.

#### 2.2.2 Hallmark-subnetwork selection

We assume the hallmarks to function independently. However, the network allows for crosstalk between hallmarks, as they can contribute to the importance of metabolites in the network which carry on to other hallmarks. To understand network characteristics on individual hallmarks, subgraphs of the MMA network were generated for each hallmark. In this process, we isolated the hallmark of interest, excluding all other hallmarks and their associated connections.

### 2.3 Node importance analysis

One form of interpretation of node importances is through the use of node centrality analysis. The usage of topology and particularly centrality measures on biological networks as a method to characterize biological mechanisms is a very well-accepted practice(Wang, Wang, and Zheng 2022). It has also been shown to provide insights into functionalities of microbial networks(Guo et al. 2022). We aim to capture information on the extent of regulation for various hallmarks by comparing their centrality scores.

With this in mind, we use eigenvector centrality as a metric of importance for nodes in the network, using the networkx nx.eigenvector_centrality function. In the case of hallmarks of aging, it gives us the influence the microbes have on the hallmarks of aging. Eigenvector centrality was chosen to account for the effect of metabolites produced by a larger number of microbes being taken to have more impact on the hallmarks as opposed to lesser produced metabolites. It does this by accounting for both the connectedness of a node, as well as how well-connected its neighboring nodes are(Negre et al. 2018). This cascading effect of influence better captures the influence of more produced metabolites on hallmarks.

For similar reasons and to provide a more thorough understanding of influence on the hallmarks, the results of eigenvector centrality were compared with Katz centrality and PageRank for the hallmarks. The networkx inbuilt nx.katz_centrality failed to converge even at high max iterations and tolerance, due to which the nx.katz_centrality_numpy approximation with attenuation factor alpha=0.1 was used with other parameters as their default. For PageRank, the inbuilt nx.pagerank with default parameters was used.

### 2.4 Node reachability analysis

Path length between nodes has been used as a method to understand the interaction between nodes in biological networks, such as in cases of metabolic genes’ shortest path lengths to disease genes being a good marker of the likelihood that the gene is a biomarker for the disease(Ni et al. 2015). In this study, path length was used as the basis to group microbes based on and identify differences in hallmarks by comparing the group sizes.

A pairwise shortest path length analysis was carried out on the directed network (unweighted), from every microbe to every hallmark. The number of microbes at a path length was counted for each hallmark. Hallmarks were then grouped based on the microbes count at path length 2 and 4. Given the number of microbes at path length two as M and number of microbes at path length 4 as N, if M/N > 2 it belongs to the first group, if M/N ∈ [2,0.5] it is in the second group and if M/N < 0.5 it is in the third group. Because we employed an unweighted network in this analysis, it essentially captures the number of metabolites or microbes required to connect a microbe with a hallmark. In reality the interactions happen at different rates i.e., have different weights. We adopted this simplification because our method’s primary objective was to categorize groups of microbes, rather than derive qualitative insights from path lengths.

### 2.5 Hallmark-microbe rank independence analysis

The MMA network captures only the nature of the association between aging hallmarks and metabolites. Reconciling the impact of different microbes on the hallmarks of aging, through their associated metabolites, is not possible without making certain simplifying assumptions (such as assuming the magnitude of impact on a given hallmark is the same for all neighboring metabolites). Alternatively, we can choose to focus our examination only on those microbes that have an unambiguous impact on a given hallmark.

#### 2.5.1 Unequivocally impacting microbes

We define a microbe B to have an unequivocal impact on a hallmark H if its impact on H is the same through every metabolite it produces/consumes. The impact of a bacteria on a hallmark through a metabolite is defined by the product of the weights on the edges between the pairs (bacteria, metabolite) and (metabolite, hallmark) in the weighted undirected network. This is measured for all microbes in MMA and microbes are removed where there exist at least 2 metabolites M1 and M2 such that the impact of the microbe on H via M1 is beneficial and via M2 is detrimental. Microbes with no association with the hallmark are also removed.

It’s possible that the strict criteria employed to define microbes with clear impact may lead to the omission of valuable information, including other microbes that have a significant influence on aging hallmarks. However, in the absence of specific research inquiries focusing on the effects of microbes or metabolites of interest on aging hallmarks, which would naturally entail relaxing the filtering parameters to include semi-ambiguous impact microbes, we deliberately opted for the most stringent definition. This decision was made to illustrate a specific use case of MMA in addressing such inquiries.

A microbe B is said to unequivocally impact aging if its impact on each hallmark is unequivocal. The MMA network yields 249 microbes that unequivocally impact aging. This gives us a 249 x 12 matrix M where M_ij_= 1, −1 or 0 if microbe i unequivocally impacts hallmark j in a beneficial manner, detrimental manner or no-effect respectively. Microbes with the same row values were grouped into modules as mapped in Supplementary Table 1. All rows of microbes from a module were removed and replaced with a representative module row. This gave us 32 modules and 74 unique rows including the modules and microbes.

#### 2.5.2 Controlling the Hallmarks of Aging

One potential therapeutics application arising from the network is the ability to regulate the hallmarks independently. A simplified solution would be to use unequivocally impacting microbes (discussed in 2.5.1). To identify the smallest set of microbes, any set of 12 linearly independent rows could be selected from the matrix M. The matrix M was converted to its row echelon form and the pivot row indices were taken using the python library sympy (the sympy.Matrix.rref() function(Meurer et al. 2017)). This generates the row echelon form matrix and returns the pivot row indices. To validate the selection, a matrix was generated with the pivot rows and the Rank of this matrix was calculated with the numpy library using the numpy.linalg.matrix_rank() function(Harris et al. 2020).

## 3 Results

### 3.1 Construction of the Microbiome-Metabolite-Aging (MMA) network

We constructed a fusion network, referred to as the Microbe-Metabolite-Aging (or MMA) network, by combining an existing extensive mammalian gut metabolic interaction network (NJC19) (Lim et al. 2020) with a novel network that captures associations between metabolites and the 12 hallmarks of aging (López-Otín et al. 2023).

We first started by fixing the definitions for each of the 12 hallmarks of aging since the specific nomenclature laid down is not standardized across literature. Next, we reconstructed the microbe-metabolite network based on NJC19 as described in Materials and Methods. The resultant 262 metabolites were used to craft PubMed queries and investigate their potential nature of association to the aging hallmarks. The extensive literature search yielded 238 unique binary associations (details are available in Supplementary Table 2). The absence of an edge between a metabolite and hallmark implies no known association to it. Finally, we combined it with the NJC19 Microbe-Metabolite network to obtain the MMA network, comprising 8418 edges and 1099 nodes. Fig.2 outlines the schematic of the curation pipeline and Table 1 describes the key topological properties of the MMA network.

**Figure 2.**
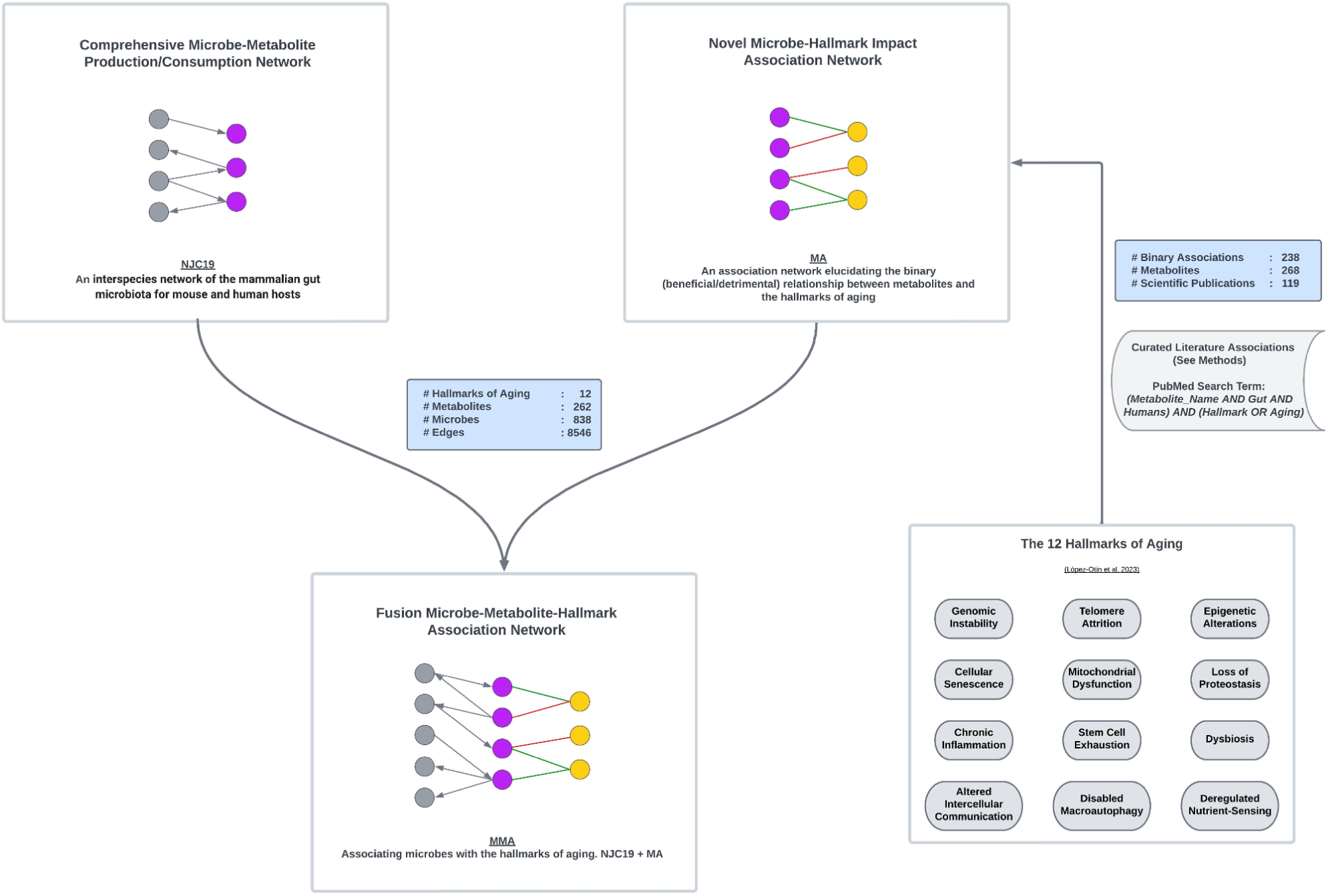
Construction of the Microbe-Metabolite-Aging (MMA) network. The flow chart of the network construction is presented. MMA is built upon literature-curated, binary associations between microbial metabolites and the 12 hallmarks of aging, combined with the extensive mammalian gut metabolic interaction network NJC19.

**Table 1.**
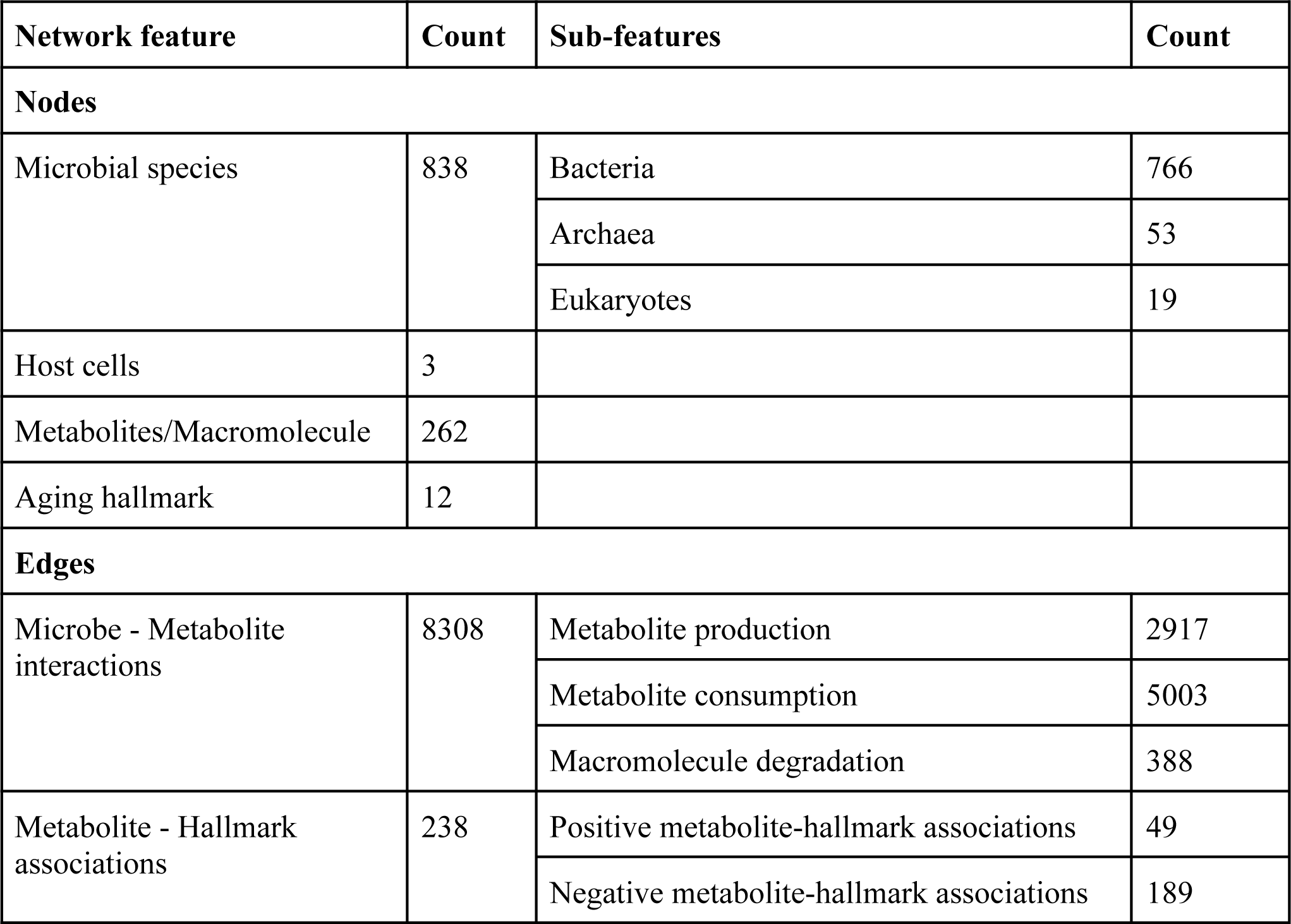
Global network statistics.

| Network feature | Count | Sub-features | Count |
| --- | --- | --- | --- |
| **Nodes** |  |  |  |
| Microbial species | 838 | Bacteria | 766 |
|  |  | Archaea | 53 |
|  |  | Eukaryotes | 19 |
| Host cells | 3 |  |  |
| Metabolites/Macromolecule | 262 |  |  |
| Aging hallmark | 12 |  |  |
| **Edges** |  |  |  |
| Microbe - Metabolite interactions | 8308 | Metabolite production | 2917 |
|  |  | Metabolite consumption | 5003 |
|  |  | Macromolecule degradation | 388 |
| Metabolite - Hallmark associations | 238 | Positive metabolite-hallmark associations | 49 |
|  |  | Negative metabolite-hallmark associations | 189 |

The undirected and unweighted MMA network is visualized in Fig.3 (the data file associated with the MMA network is available in Supplementary Network 1). Due to the high number of nodes resulting in an unreadable dense graph, nodes of the same type are merged for visualization. Fig.4 shows a sub-network of Fig.3, specifically the metabolite-aging hallmark network (depicted by the orange and yellow nodes respectively in Fig.3).

**Figure 3.**
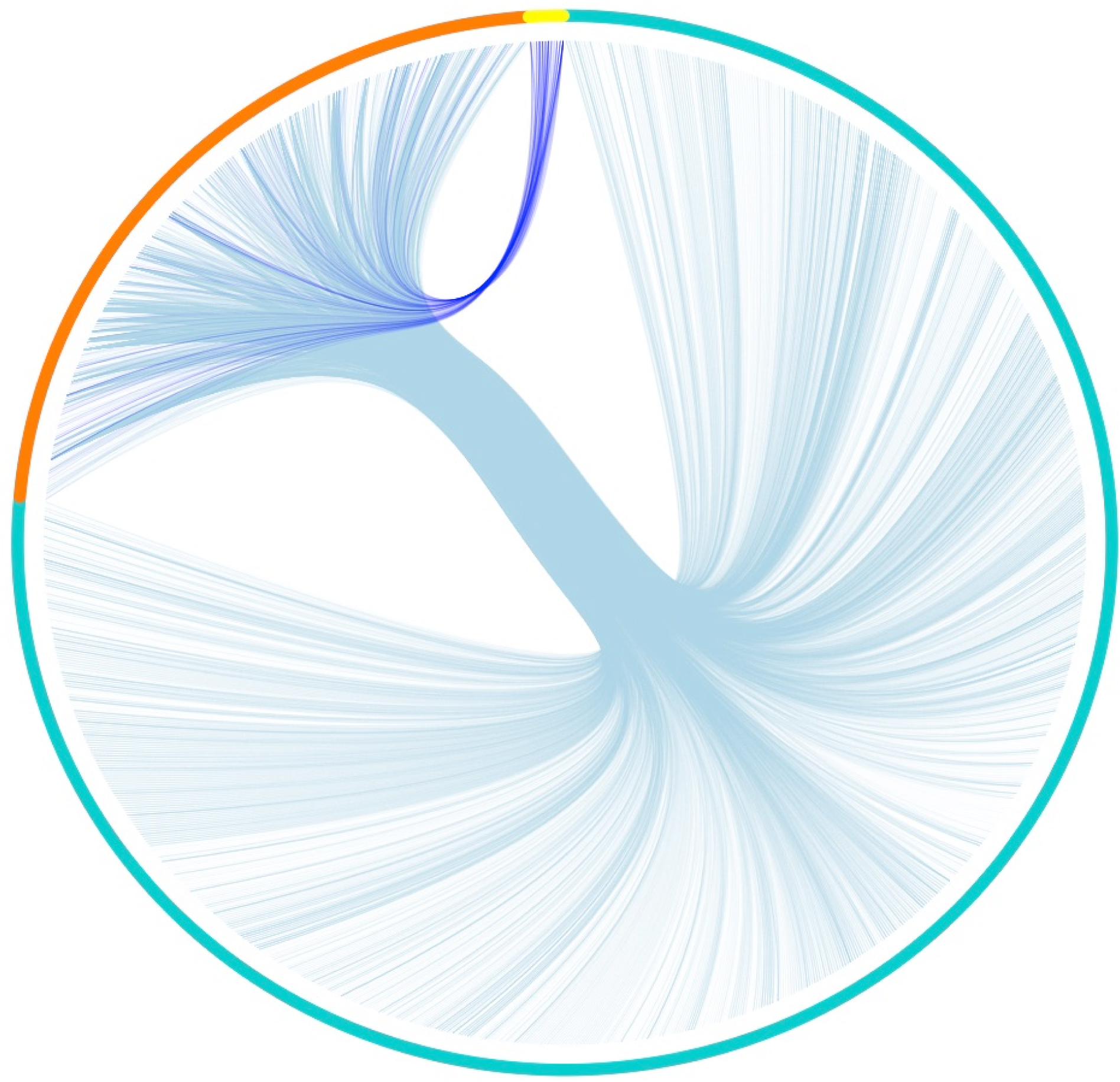
Chord diagram depicting the MMA network, the outer ring depicts the nodes and the arcs inside are edges. Blue: Microbes, Orange: Metabolites, Yellow: Hallmarks. Light Blue: Microbe-Metabolite Edges, Dark Blue: Metabolite-Hallmark Edges.

**Figure 4.**
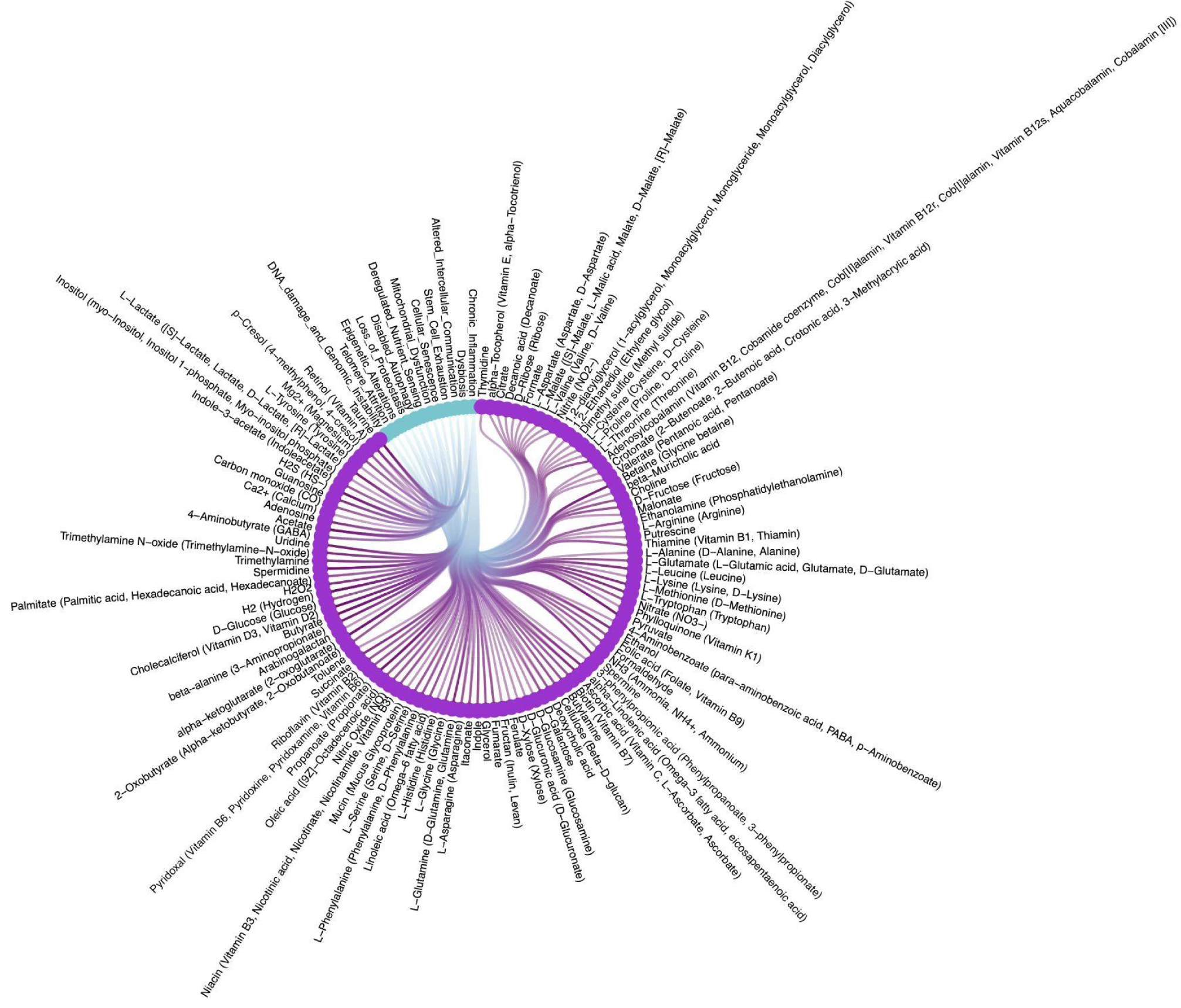
Chord diagram depicting the Metabolite-Aging Hallmark network. Blue nodes: Hallmarks, Violet nodes: Metabolites, arcs in between are unweighted edges from the metabolite to the hallmarks.

The MMA network serves as a tool to investigate the mammalian (and in particular human) gut microbiome as an effector of aging in a systems-level manner.

### 3.2 Differential regulation of aging hallmarks by microbial metabolites

The MMA network provides a foundational knowledge graph to study the dynamics and impact of microbes and their metabolites on the different hallmarks of aging. While various studies [(Imanikia et al. 2019), (Zhou, Hu, and Wang 2021; Shmookler Reis et al. 2011; Mansfeld et al. 2015)(Zhou, Hu, and Wang 2021; Shmookler Reis et al. 2011; Ramachandran et al. 2019)] have outlined the role of microbial metabolites in regulating specific hallmarks of aging, the MMA network allows a systematic analysis of the differential regulation of these hallmarks by microbial metabolites based on the topological properties of the network.

Fig.5a shows the metabolite association distribution for the hallmarks of aging in the MMA network where the number of metabolites is represented in the Y-axis against the 12 hallmarks in the X-axis. As elucidated in the figure, the hallmarks have distinctly different metabolite associations. While chronic inflammation, mitochondrial dysfunction, and loss of proteostasis have the largest number of metabolites associated with each of them in the MMA, the metabolite-mediated regulation has less impact on the hallmarks of Telomere attrition and stem cell exhaustion. The results suggest the role of microbes in mediating aging through inflammation and metabolism axes of aging.

**Figure 5a.**
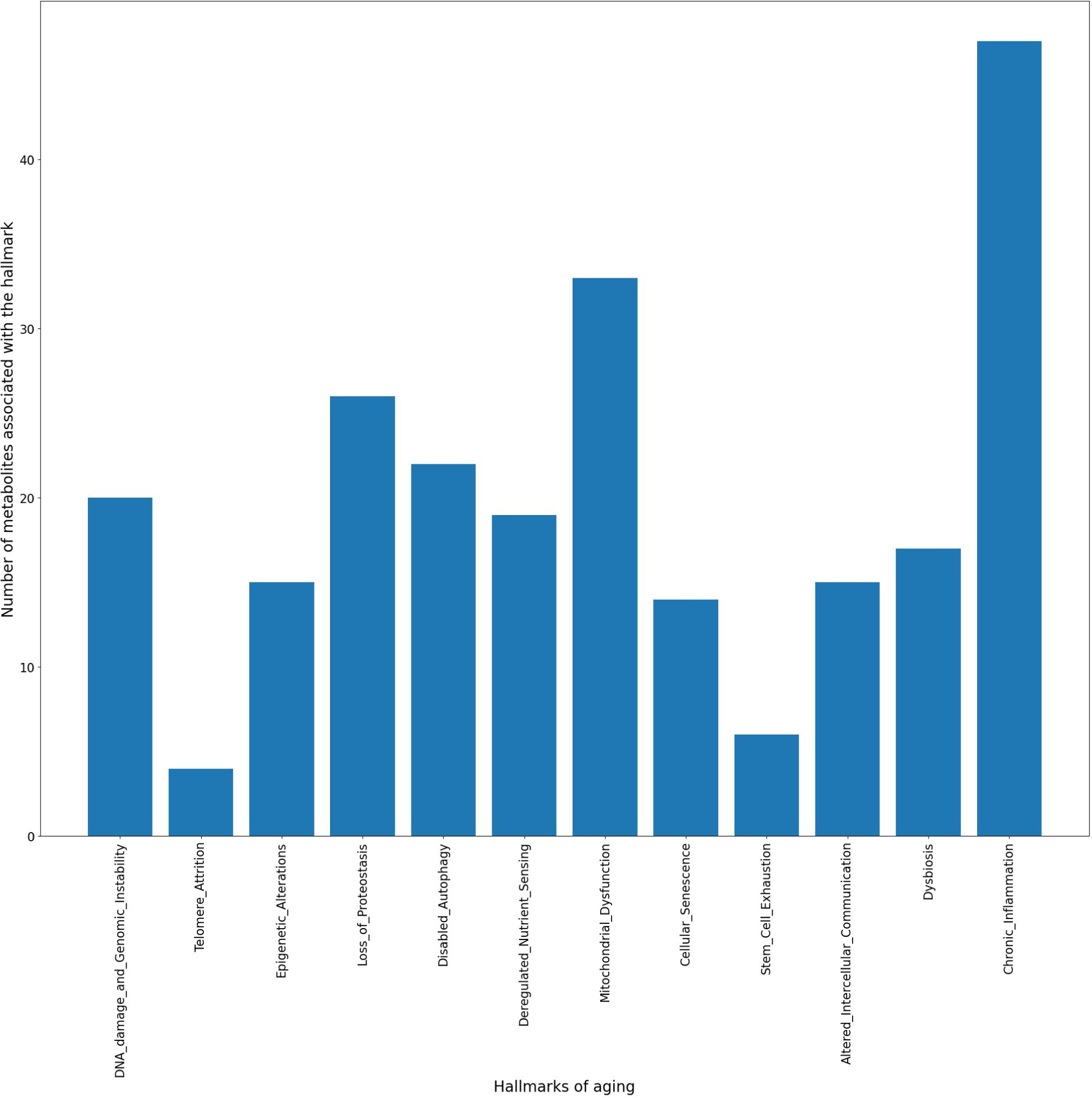
Metabolite association distribution for the hallmarks of aging in the MMA network where the number of metabolites is represented in the Y-axis against the 12 hallmarks in the X-axis.

While the number of metabolites associated with a hallmark gives an aggregate measure of their role, the influence of some metabolites depend on their connectivity to other metabolites and the role of those metabolites in regulating aging hallmarks in the network. Fig. 5b, compares the hallmarks based on their influence score (measured by Eigenvector centrality, the details of which are outlined in Materials and Methods). As seen from the plot, while mitochondrial dysfunction and chronic inflammation have higher eigenvector centrality (similar to the higher metabolites associated reported earlier), other hallmarks, particularly genomic instability as well as deregulated nutrient sensing emerge as high influencers (high Eigenvector centrality scores) in the MMA network. In the case of mitochondrial dysfunction, the association with formate and fructose (which are not associated with genomic instability, nutrient sensing, or chronic inflammation, but have a high degree in the network, i.e., are produced, consumed, or regulated by more bacteria) raise the eigenvector centrality. However, in the case of genomic instability, the influence score (eigenvector centrality) is affected by connectivity to ethanol which pushes the score up and brings the influence of genomic instability to a similar level as that of mitochondrial dysfunction. Furthermore, influence was also estimated by other transitive centrality metrics, Katz centrality and PageRank, as depicted in Fig. 5c and 5d respectively, and the results compared with eigenvector centrality. Katz centrality showed similar results to eigenvector centrality, while the PageRank is notably lower (when comparing rank order in the centrality measure) for epigenetic alterations and nutrient sensing, as well as higher for cellular senescence and stem cell exhaustion. Implying the possibility of more complex transitive influence of microbes and metabolites on the hallmarks, such as the likelihood of metabolites ending up performing a role associated with the hallmarks as opposed to being consumed by microbes, as is captured by PageRank.

**Figure 5b.**
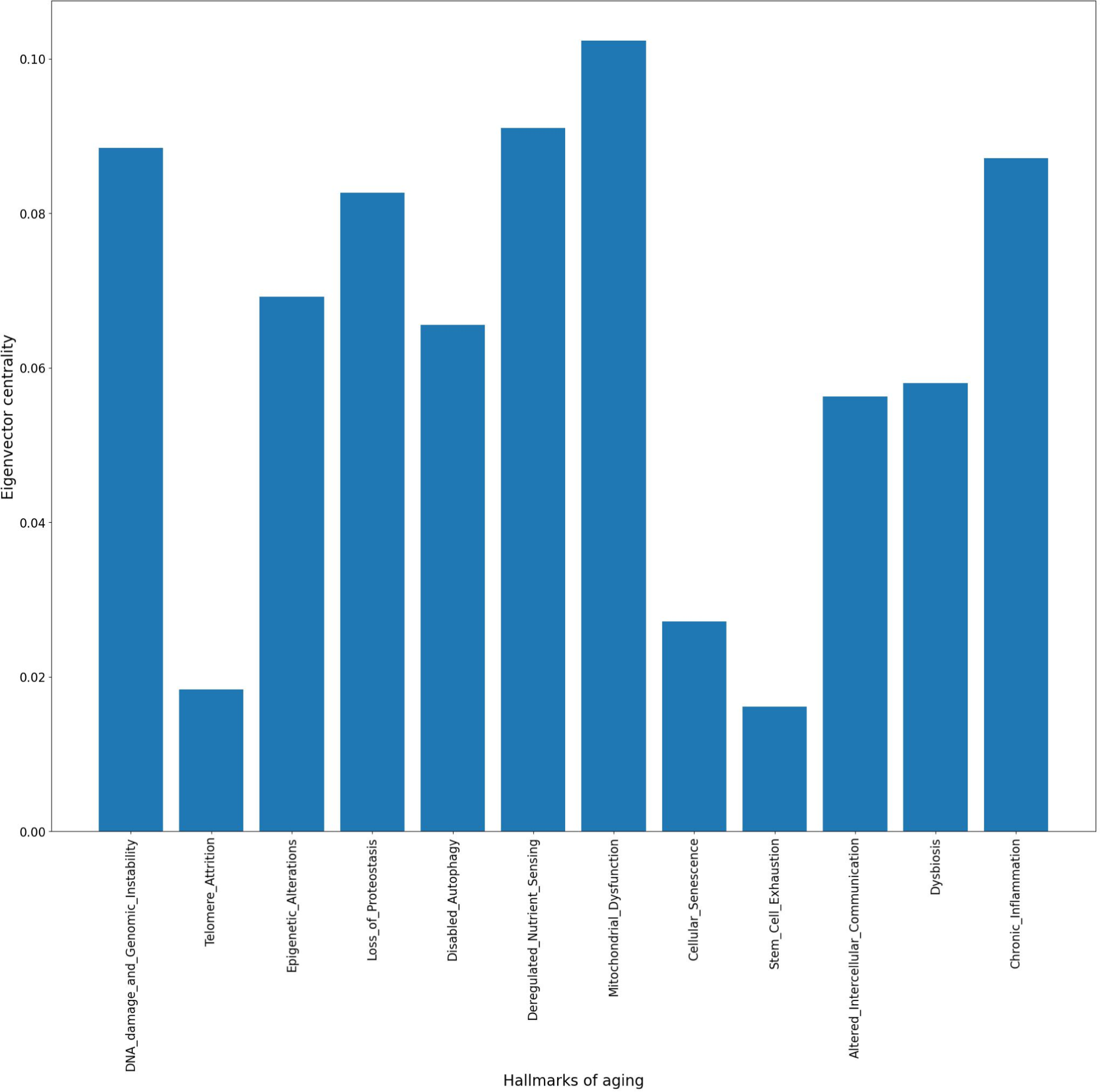
Network influence on the hallmarks, captured through the eigenvector centrality distribution

**Figure 5c.**
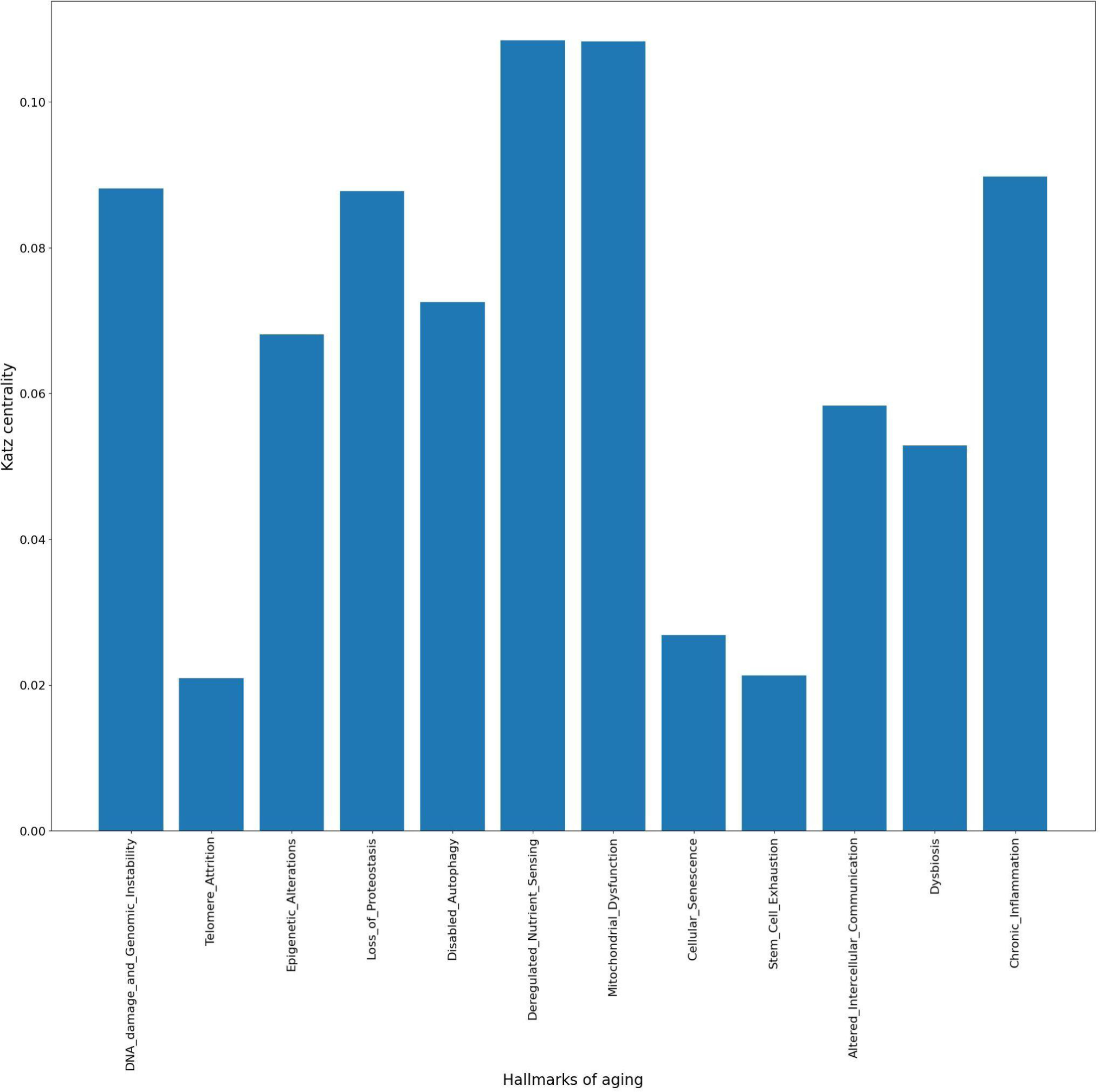
Network influence on the hallmarks, captured through the Katz centrality distribution

**Figure 5d.**
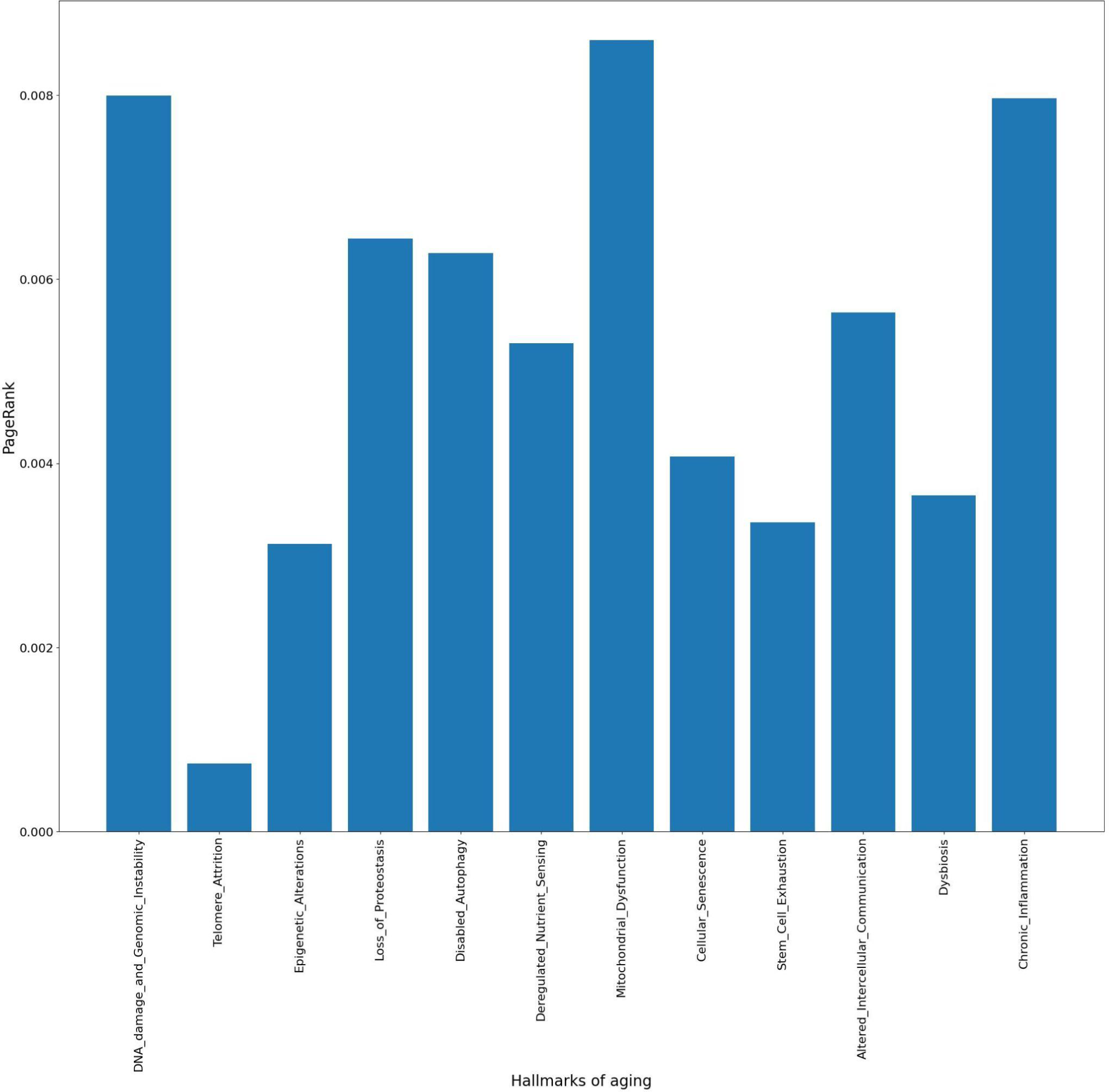
Network influence on the hallmarks, captured through the PageRank distribution

These analyses based on the MMA network model help to identify the differential regulatory role of microbial metabolites on the hallmarks of aging, both at the aggregate association levels as well as at the topological properties of the individual metabolites and their influence on the network.

### 3.3 Identification of core microbes for the hallmarks of aging

The path-length or hop length of the shortest path connecting nodes is a metric used often in analysis of biological networks (Ren, Ay, and Kahveci 2018). It has also seen usage in the context of identifying the genetic determinants of aging (Managbanag et al. 2008). For the purpose of this analysis, the directed version of the network was used to better characterize the existence and lengths of paths. Fig.6 shows the number of microbes at different path lengths from each of the hallmarks. As seen in the figure, majority of the hallmarks have between 460-550 microbes at path length 2 (i.e. core microbes for the hallmark - microbes producing metabolites directly associated with the hallmark), 90-180 microbes at a path length 4, i.e. microbes which produce metabolites that are then consumed by the core microbes, and 10-25 microbes that are at a path length of 6 which produce metabolites consumed by microbes at a path length of 4. Segregation of microbes using their path length to a hallmark as a metric serves as a method to cluster microbes based on the mode and the extent to which they impact the hallmarks.

**Figure 6.**
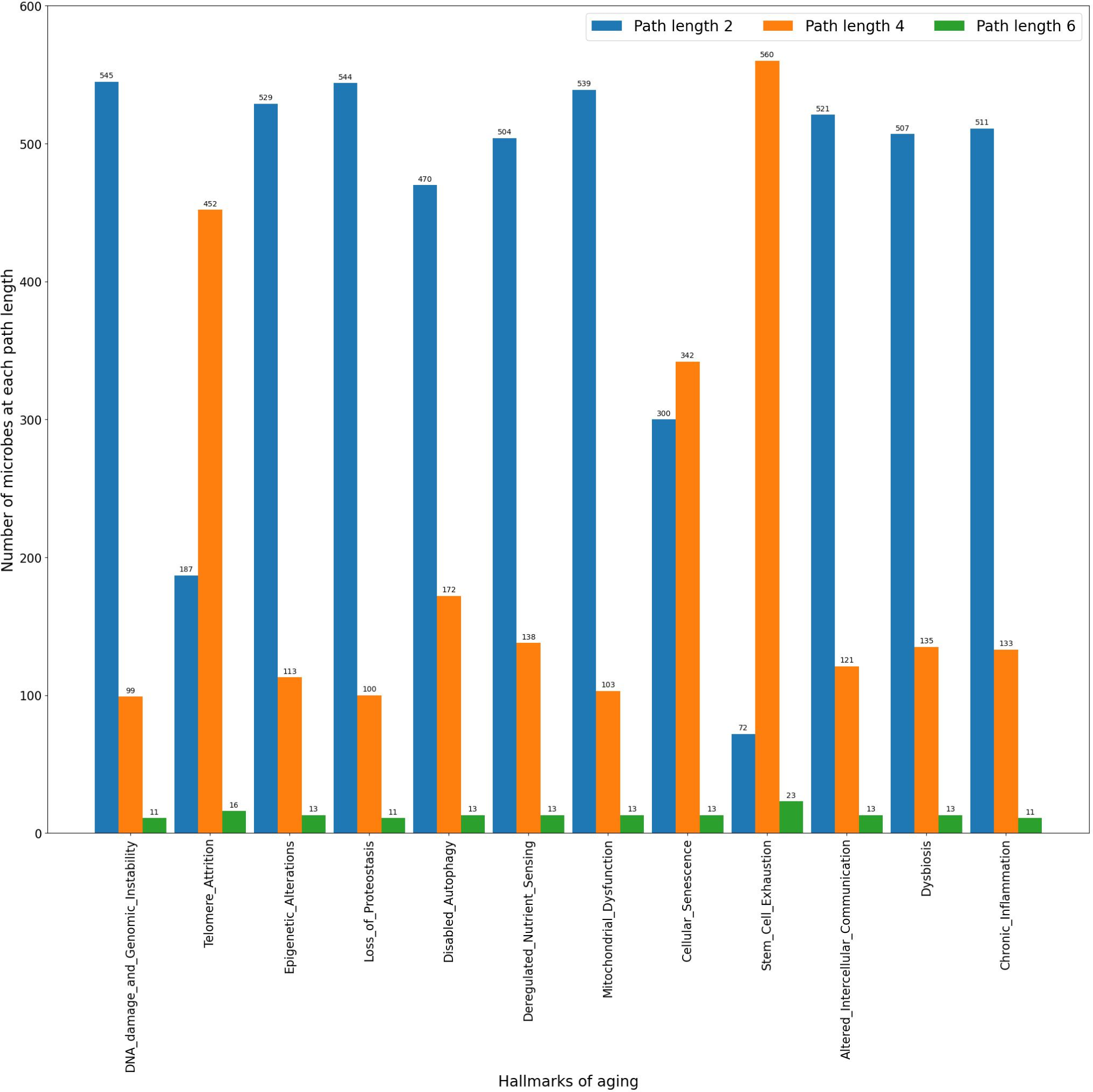
Hallmarks of Aging and the number of Microbes with a directed path from the microbe to the Hallmark. Blue, Orange and Green correspond to the number of microbes with a path length of 2, 4 and 6 respectively. There exists a metabolite between every link from the hallmark to the microbe as well as from a microbe to another microbe, hence the path lengths are in multiples of two.

While there are reasons for predicting a relationship between path length and functional specificity of nodes in biological networks (Xu et al. 2011), the usage of path length has limitations in the ability to capture the complex nature of biological systems, where the biological significance can depend on specific context, types of nodes and interactions. In this context, our analysis is focused on clustering of microbes based on their impact on a hallmark through their metabolites and not on the specific impact of a particular path-length on the mechanism of impact.

In Fig.6, we observe hallmarks with different relationships between the number of microbes at path length 4 and those at path length 2. This also gives rise to three broad classes of hallmarks; the first class consists of majority of hallmarks, where there are significantly more core microbes than microbes at path length 4, the second class consists of hallmarks where the core microbes and the microbes at path length 4 are comparable to each other; and the third class where the core microbes are significantly less than those at path length 4 (the exact ranges for the classes are defined in Materials and Methods). Based on this analysis, genomic instability, epigenetic alterations, loss of proteostasis, disabled autophagy, deregulated nutrient sensing, mitochondrial dysfunction, altered intercellular communication, dysbiosis and chronic inflammation come under the first class. The second class consists of the hallmark cellular senescence while the last class contains hallmarks telomere attrition and stem cell exhaustion.

The formation of classes within hallmarks based on the varying impact of microbes as identified from their topological properties in the network (path length) provides unique insights into the relative biological role on the hallmark. This implies an importance of the network topology in granting characteristics to biological aging mediated by the microbiome.

### 3.4 Characterization of Microbial regulation of the Hallmarks of Aging

The MMA graph contains no direct association between the Microbes and the Hallmarks of Aging as the association is mediated through intermediate metabolites. In order to characterize the direct impact of the microbes on the hallmarks, the network was linearly transformed from a tripartite graph of the microbes-metabolites-hallmark to a microbe-hallmark network. Further, it was filtered for contradictory effects, as described in the Materials and Methods (the final adjacency matrix and pairwise interaction file are in Supplementary Network 4 and 5 respectively). The microbe-hallmark network focused on microbes which had a defined, directional effect on a hallmark, referred to as the unequivocal microbe-hallmark network, by the removal of microbes which had contradictory effects on a hallmark due to different interactions with metabolites.

The network contains 249 microbes with an unequivocal effect on each hallmark (details are available in Supplementary Network 4). Subsequently, microbes were clustered based on their impact on Aging as a whole, i.e., microbes with the same impact on all 12 hallmarks were grouped into a module (module details are available in Supplementary Table 1).

This analysis identified 74 unique modules of modes of action on the hallmarks of aging as seen in Fig.7. On this network linear independence analysis to identify if it was possible to control each of the 12 hallmarks independently through regulation of the unequivocal microbes. The rank of the adjacency matrix for the network is 12, equal to the number of hallmarks, implying that every hallmark of aging can be controlled through the increase or decrease of microbes in these modules.

**Figure 7.**
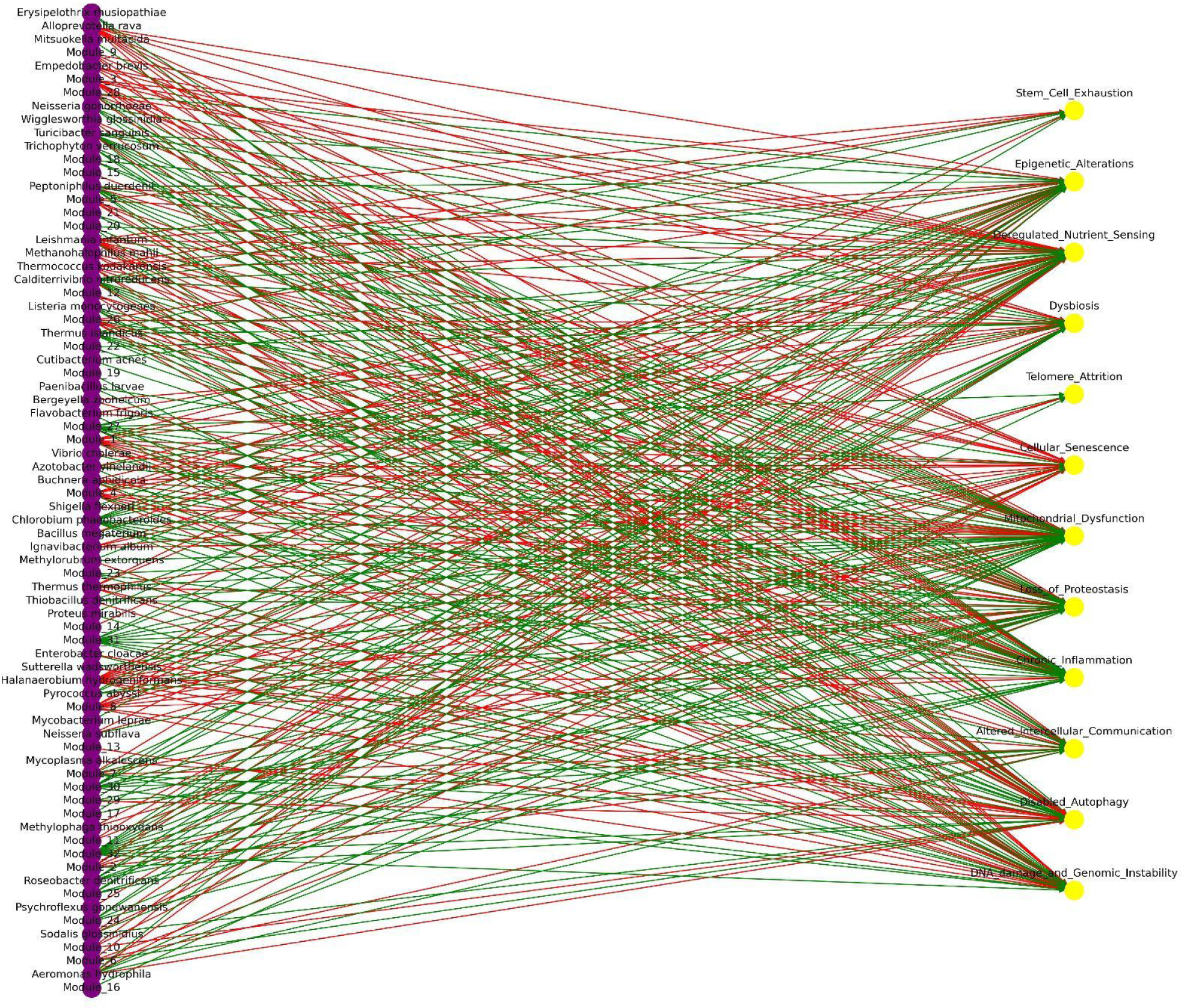
Bipartite plot of microbe/microbial module unequivocal association with aging hallmarks. Yellow nodes: Aging hallmarks, Purple nodes: Microbes or microbe modules. Green edges: Positive associations, Red edges: Negative associations.

We investigated the smallest community of microbes that can be used to differentially affect each hallmark of aging (either positively, negatively, or having no-impact) by making the simplifying assumption that each microbe can be independently controlled.

We identified a subset of the 74 microbes and modules that are linearly independent on their impact on the hallmarks and can hence regulate any single hallmark through its own regulation as outlined in the Materials and Methods. The identified set contains the following 12 microbes: Halanaerobium hydrogeniformans, Alloprevotella rava, Leishmania infantum, Melissococcus plutonius(Module_1), Leptotrichia goodfellowii(Module_2), Prevotella veroralis(Module_3), Caldisericum exile(Module_5), Lactobacillus ingluviei(Module_6), Leuconostoc kimchii(Module_7), Ruminococcus lactaris(Module_8), Sphingobium japonicum(Module_9), Empedobacter brevis. It may be noted that in order to list the minimal set of microbes differentially impacting each hallmark of aging, microbes were chosen arbitrarily from a selected module as all microbes are treated equal in any given module. It is possible in future analysis to consider microbes from modules which may be selected based on prior knowledge of ability to independently control the microbes, such as specific microbes that consume certain metabolites, or are susceptible to specific narrow-spectrum antibiotics.

## 4 Discussion

We introduce a fusion network (MMA) that combines an extensive mammalian metabolic interaction network (Lim et al. 2020) with a novel network borne out of extensive manual literature review, that associates the network metabolites with the hallmarks of aging. We analyze MMA to provide insights into the regulatory mechanisms underlying biological aging and position this network as a comprehensive ground truth for future modeling studies and mechanistic hypothesis generation and discovery engines.

We constructed a fusion network model of microbes, metabolites and hallmarks of aging with the goal to collect, curate and harmonize the current state of art knowledge on the complex interaction dynamics of these components. The large-scale MMA network provides a model to investigate the complex interplay of microbes and their metabolites as effectors of aging hallmarks at a systems level. At the same time, it is pertinent to elucidate the boundaries of the MMA network. In particular, it is to be noted, as outlined in the previous sections, that the interactions captured are primarily based on associations from primary research and not specifically causative or mechanistic underpinnings. The MMA network construction protocol incorporated steps to avoid ambiguities in the literature by taking the context of the underlying biology into account where applicable, looking for the research to be applicable in the context of aging or similar accumulative effects.

Our network accounts for microbial interactions by competition, mutualism through their interactions with common metabolites, while in reality, interactions between microbes go beyond metabolite-mediated interactions and include complex interactions such as quorum sensing and cellular transduction signals. Similarly, some metabolites may impact aging only in the presence of other metabolites, leading to more complex dynamics than linear associations.

The MMA protocol captures the direct effect of metabolites on hallmarks independently. However, hallmarks are known to have complex systems-level interactions with others, such as feedback loops, etc (Cohen et al. 2020). Further research can incorporate the complex nature of these associations, which may be modeled by the addition of transformation layers in between the microbe and metabolite layer as well as the metabolite and aging hallmark layer, along with edges between the nodes in each layer.

Real-world MMA networks will be subsets of the global network, with edges from microbes possibly being weighted by their abundance in the gut. The results of the analyses applied to these networks and the subsequent comparison to clinical observations will allow for the validation of the network. The usage of topological analysis alone may not capture aspects of these interactions such as their dynamic nature, which can be influenced by host factors as well as the environment. However, further testing is needed to determine the scope of this influence.

The MMA network can serve as a modeling tool for a variety of scientific hypotheses generation. This work focused analysis on studying the effect of microbes and metabolites on the 12 hallmarks of aging. Network properties like association distribution and influence measures like eigenvector centrality, katz centrality and pagerank identified the differential regulatory role of microbial metabolites on the hallmarks of aging, specifically mitochondrial dysfunction, chronic inflammation, genomic instability and deregulated nutrient sensing. Connectivity analysis based on path-length from microbes to hallmarks (mediated through their metabolites) classified the hallmarks into three major groups based on the role of core, secondary and tertiary microbes on them. Further, on network transformation wherein the direct connection from microbes to hallmarks were identified based on directional effect, the unequivocal microbe-hallmark network allowed the characterization of microbes based on their unequivocal, i.e defined and directional effect on each hallmark.

The above analyses elucidate the complex mechanisms underlying the dynamics of microbial metabolite mediated effect on aging through its hallmarks. As outlined in this work, the biology of the interactions, together with their topological properties in the larger context of the network play a crucial role in differentiating aging hallmarks through the lens of the microbiome. We highlight how the MMA network model can be further developed, when paired with mathematical modeling techniques, in future applications to unravel the dynamics of aging mediated through the microbiome.

### 4.1 Mechanistic Hypothesis Generation and Discovery

Microbiome science, in order to continue growing the way it has been, will benefit from a focus on causation and mechanism (Fischbach 2018). The field is moving towards measurable and unambiguous phenotype-microbiome associations. Generating and testing such hypotheses in vivo is time consuming and demands intelligent experimental design. Along with assisting hypothesis generation in the area of aging, the MMA network can also assist metabolomic investigation using metagenomic data (16S and WMGS) by focusing on the corresponding aging microbial drivers.

Alternatively, the metabolite-aging network can be paired along with gut metabolomics data that provides in vivo information about the metabolite concentrations in the gut to identify metabolites of interest based on the magnitude of impact on the hallmarks of aging.

### 4.2 Optimal Communities for Healthy Aging

Recent advancements in microbial community modeling through Genome Scale Metabolic Models (GMMs) has opened up the possibility of modeling metabolite fluxes in the gut microbiome using metagenomic data (Diener, Gibbons, and Resendis-Antonio 2020). Paired with GMMs of host cells, one can model host-microbe interactions. Such analyses open the possibility of modeling organ-microbiome metabolite exchange and serve as a virtual diagnostic tool for mechanistic hypothesis generation (Thiele et al. 2020).

The MMA network can serve as a tool to measure (track and compare) the impact of a given microbial community on healthy aging. It is worth noting that the MMA network primarily enables qualitative insights into the microbiome. GMMs focus on the stoichiometry of microbial biochemistry, yielding highly quantitative outputs regarding metabolic fluxes and microbial growth rates. However, these quantitative outcomes are subject to sensitivity based on input parameters and the representation of host-microbe relationships within the simulated environment. Requiring precise definitions of the simulated environment, which due to the difficulty of measuring is a potential source for errors.

However, the MMA network can also be used to craft aging specific objective functions (or constraints) that maximize host benefit to healthy aging. This could be used as a simple extension to pre-existing GMM based tools that are used to identify the minimal direct interventions to a microbial community that results in the increased (or maximal) production of a specific compound (Zomorrodi and Maranas 2012).

Alternatively, one could use the framework of GMM’s (Zomorrodi and Maranas 2012; Diener, Gibbons, and Resendis-Antonio 2020) on microbial communities along with in-silico optimization to identify theoretical chemical cocktails which can be administered as drugs, nutraceuticals or simply through nutrition, that could drive concentrations of key metabolites identified in the MMA network. This will allow for targeted clinical trial design to accelerate the discovery of chemical modulators of healthy aging.

Aging is a complex, multifactorial process which is governed by interaction of multiple processes at different levels. The microbiome, through its metabolites plays an important role in modulating aging, particularly the hallmark processes associated with aging. It may be noted that the impact of a given microbial community on the individual hallmarks of aging can allow stratifying microbiome datasets based on their mode of action, leading to the potential of using the gut microbiome as a precision medicine tool for therapeutic stratification (Schupack et al. 2022) in healthy aging. The MMA network model elucidates how all hallmarks are not equal from the lens of the microbiomes. Thus, studying the effect of microbiome on aging requires robust computational models together with novel experimental design and validation. We envisage the MMA network presented in this work to serve as a foundational model towards deeper understanding of aging mediated through the microbiome.

## Supporting information

Microbes to Module mapping

Data on metabolite and hallmark associations along with reference publications

Undirected and unweighted network

Undirected and weighted network

Directed and weighted network

Final adjacency matrix

Pairwise interaction network

## Supplementary Material

Supplementary Table 1.csv : Microbes to Module mapping

Supplementary Table 2.csv: Data on metabolite and hallmark associations along with reference publications

Supplementary Network 1.csv: Undirected and unweighted network

Supplementary Network 2.csv: Undirected and weighted network

Supplementary Network 3.csv: Directed and weighted network

Supplementary Network 4.csv: Final adjacency matrix

Supplementary Network 5.csv: Pairwise interaction network

## Notes

### Competing Interest Statement

SM, NBD and BPK are employed by Iom Bioworks. BPK and SG are directors of Iom Bioworks with equity interests.

### Summary of Updates

Motivations added to individual sections in the methods; Added Katz centrality and PageRank for measurement of influence of microbes on hallmarks; Possible shortcomings of the methods and analysis were also added

## References

Agus, Allison, Karine Clément, and Harry Sokol. 2021. “Gut Microbiota-Derived Metabolites as Central Regulators in Metabolic Disorders.” Gut 70 (6): 1174–82.

Badal, Varsha D., Eleonora D. Vaccariello, Emily R. Murray, Kasey E. Yu, Rob Knight, Dilip V. Jeste, and Tanya T. Nguyen. 2020. “The Gut Microbiome, Aging, and Longevity: A Systematic Review.” Nutrients 12 (12). 10.3390/nu12123759.

Cohen, A. A., Ferrucci, L., Fülöp, T., Gravel, D., Hao, N., Kriete, A., Levine, M. E., Lipsitz, L. A., Olde Rikkert, M. G. M., Rutenberg, A., Stroustrup, N., & Varadhan, R. 2022. “A complex systems approach to aging biology.” Nature aging 2(7): 580–591.

Diener, Christian, Sean M. Gibbons, and Osbaldo Resendis-Antonio. 2020. “MICOM: Metagenome-Scale Modeling To Infer Metabolic Interactions in the Gut Microbiota.” mSystems 5 (1).

Fischbach, Michael A. 2018. “Microbiome: Focus on Causation and Mechanism.” Cell 174 (4): 785–90.

Ghosh, Tarini Shankar, Fergus Shanahan, and Paul W. O’Toole. 2022. “The Gut Microbiome as a Modulator of Healthy Ageing.” Nature Reviews. Gastroenterology & Hepatology 19 (9): 565–84.

Guo, Bing, Lei Zhang, Huijuan Sun, Mengjiao Gao, Najiaowa Yu, Qianyi Zhang, Anqi Mou, and Yang Liu. 2022. “Microbial Co-Occurrence Network Topological Properties Link with Reactor Parameters and Reveal Importance of Low-Abundance Genera.” Npj Biofilms and Microbiomes 8 (1): 1–13.

Hagberg, Aric, Pieter Swart, and Daniel S Chult. 2008. “Exploring Network Structure, Dynamics, and Function Using Networkx.” LA-UR-08-05495; LA-UR-08-5495. Los Alamos National Lab. (LANL), Los Alamos, NM (United States). https://www.osti.gov/servlets/purl/960616.

Harris, Charles R., K. Jarrod Millman, Stéfan J. van der Walt, Ralf Gommers, Pauli Virtanen, David Cournapeau, Eric Wieser, et al. 2020. “Array Programming with NumPy.” Nature 585 (7825): 357–62.

Imanikia, Soudabeh, Ming Sheng, Cecilia Castro, Julian L. Griffin, and Rebecca C. Taylor. 2019. “XBP-1 Remodels Lipid Metabolism to Extend Longevity.” Cell Reports 28 (3): 581–89.e4.

Kim, Minhoo, and Bérénice A. Benayoun. 2020. “The Microbiome: An Emerging Key Player in Aging and Longevity.” Translational Medicine of Aging 4 (July): 103–16.

Kowald, A., and T. B. Kirkwood. 1996. “A Network Theory of Ageing: The Interactions of Defective Mitochondria, Aberrant Proteins, Free Radicals and Scavengers in the Ageing Process.” Mutation Research 316 (5-6): 209–36.

Lim, Roktaek, Josephine Jill T. Cabatbat, Thomas L. P. Martin, Haneul Kim, Seunghyeon Kim, Jaeyun Sung, Cheol-Min Ghim, and Pan-Jun Kim. 2020. “Large-Scale Metabolic Interaction Network of the Mouse and Human Gut Microbiota.” Scientific Data 7 (1): 204.

López-Otín, Carlos, Maria A. Blasco, Linda Partridge, Manuel Serrano, and Guido Kroemer. 2013. “The Hallmarks of Aging.” Cell 153 (6): 1194–1217.

López-Otín, Carlos, Maria A. Blasco, Linda Partridge, Manuel Serrano, and Guido Kroemer. 2023. “Hallmarks of Aging: An Expanding Universe.” Cell 186 (2): 243–78.

Managbanag, J. R., Tarynn M. Witten, Danail Bonchev, Lindsay A. Fox, Mitsuhiro Tsuchiya, Brian K. Kennedy, and Matt Kaeberlein. 2008. “Shortest-Path Network Analysis Is a Useful Approach toward Identifying Genetic Determinants of Longevity.” PloS One 3 (11): e3802.

Mansfeld, Johannes, Nadine Urban, Steffen Priebe, Marco Groth, Christiane Frahm, Nils Hartmann, Juliane Gebauer, et al. 2015. “Branched-Chain Amino Acid Catabolism Is a Conserved Regulator of Physiological Ageing.” Nature Communications 6 (December): 10043.

Martin, Alyce M., Emily W. Sun, Geraint B. Rogers, and Damien J. Keating. 2019. “The Influence of the Gut Microbiome on Host Metabolism Through the Regulation of Gut Hormone Release.” Frontiers in Physiology 10 (April): 428.

Meurer, Aaron, Christopher P. Smith, Mateusz Paprocki, Ondřej Čertík, Sergey B. Kirpichev, Matthew Rocklin, Amit Kumar, et al. 2017. “SymPy: Symbolic Computing in Python.” PeerJ Computer Science 3 (January): e103.

Negre, Christian F. A., Uriel N. Morzan, Heidi P. Hendrickson, Rhitankar Pal, George P. Lisi, J. Patrick Loria, Ivan Rivalta, Junming Ho, and Victor S. Batista. 2018. “Eigenvector Centrality for Characterization of Protein Allosteric Pathways.” Proceedings of the National Academy of Sciences of the United States of America 115 (52): E12201–8.

Ni, Qi, Xianming Su, Jingqi Chen, and Weidong Tian. 2015. “Prediction of Metabolic Gene Biomarkers for Neurodegenerative Disease by an Integrated Network-Based Approach.” BioMed Research International 2015 (May): 432012.

Ragonnaud, Emeline, and Arya Biragyn. 2021. “Gut Microbiota as the Key Controllers of ‘Healthy’ Aging of Elderly People.” Immunity & Ageing: I & A 18 (1): 2.

Rajilić-Stojanović, Mirjana, and Willem M. de Vos. 2014. “The First 1000 Cultured Species of the Human Gastrointestinal Microbiota.” FEMS Microbiology Reviews 38 (5): 996–1047.

Ramachandran, Prasanna V., Marzia Savini, Andrew K. Folick, Kuang Hu, Ruchi Masand, Brett H. Graham, and Meng C. Wang. 2019. “Lysosomal Signaling Promotes Longevity by Adjusting mitochondrial Activity.” Developmental Cell 48 (5): 685–96.e5.

Ren, Yuanfang, Ahmet Ay, and Tamer Kahveci. 2018. “Shortest Path Counting in Probabilistic Biological Networks.” BMC Bioinformatics 19 (1): 465.

Schupack, Daniel A., Ruben A. T. Mars, Dayne H. Voelker, Jithma P. Abeykoon, and Purna C. Kashyap. 2022. “The Promise of the Gut Microbiome as Part of Individualized Treatment Strategies.” Nature Reviews. Gastroenterology & Hepatology 19 (1): 7–25.

Sharma, Rishi, and Arvind Ramanathan. 2020. “The Aging Metabolome-Biomarkers to Hub Metabolites.” Proteomics 20 (5-6): e1800407.

Shmookler Reis, Robert J., Lulu Xu, Hoonyong Lee, Minho Chae, John J. Thaden, Puneet Bharill, Cagdas Tazearslan, et al. 2011. “Modulation of Lipid Biosynthesis Contributes to Stress Resistance and Longevity of C. Elegans Mutants.” Aging 3 (2): 125–47.

Sung, Jaeyun, Seunghyeon Kim, Josephine Jill T. Cabatbat, Sungho Jang, Yong-Su Jin, Gyoo Yeol Jung, Nicholas Chia, and Pan-Jun Kim. 2017. “Global Metabolic Interaction Network of the Human Gut Microbiota for Context-Specific Community-Scale Analysis.” Nature Communications 8 (June): 15393.

Thiele, Ines, Swagatika Sahoo, Almut Heinken, Johannes Hertel, Laurent Heirendt, Maike K. Aurich, and Ronan Mt Fleming. 2020. “Personalized Whole-Body Models Integrate Metabolism, Physiology, and the Gut Microbiome.” Molecular Systems Biology 16 (5): e8982.

Wang, Mengyuan, Haiying Wang, and Huiru Zheng. 2022. “A Mini Review of Node Centrality Metrics in Biological Networks.” International Journal of Network Dynamics and Intelligence, December, 99–110.

Xu K, Bezakova I, Bunimovich L, Yi SV. 2011. “Path lengths in protein-protein interaction networks and biological complexity.” Proteomics 11 (10): 1857–67.

Zhou, Yue, Guo Hu, and Meng C. Wang. 2021. “Host and Microbiota Metabolic Signals in Aging and Longevity.” Nature Chemical Biology 17 (10): 1027–36.

Zomorrodi, Ali R., and Costas D. Maranas. 2012. “OptCom: A Multi-Level Optimization Framework for the Metabolic Modeling and Analysis of Microbial Communities.” PLoS Computational Biology 8 (2): e1002363.

